# The 3D point spread function can be autoencoded in latent space

**DOI:** 10.1101/2024.10.14.618231

**Authors:** Axel Truedson, Konrad Gras, Johan Elf

**Affiliations:** Department of Cell and Molecular Biology, SciLifeLab, Uppsala University

## Abstract

The 3D point spread function of a fluorescence emitter in a living cell is often different from that of the z-stack of bead images typically used as a reference. Here we show that the location in z of a fluorescent emitter can be directly accessed in the latent space of an autoencoder trained on the sample images without z-reference data. The corresponding decoding network represents an accurate point spread function.

---

Point spread function (PSF) engineering is commonly used in single-molecule localization microscopy to access 3D information^1,2^. Modifications in the optical path are introduced to make point emitters at different z-depth produce different image patterns. By comparing the pattern from an emitter to a reference 3D PSF, it is possible to estimate the z position of the point emitter. The reference 3D-PSF is typically determined by obtaining a z-stack of bead images and fitting the image stack to an interpolated 3D function^3^. An example of such an image stack taken through an astigmatic lens at different z positions is shown in Fig 1A. The drawback of this method is that bead images rarely accurately represent the single molecule emitters in living cells due to differences in spectra, size, aberrations introduced by the sample, and signal to background properties^4^. It is therefore desirable to determine 3D-PSFs directly for the emitters of interest in the sample. This has been problematic since z-stacks of real emitters cannot be obtained due to bleaching or, in the case of live cell experiments, because the emitter is moving.

**Figure 1.**
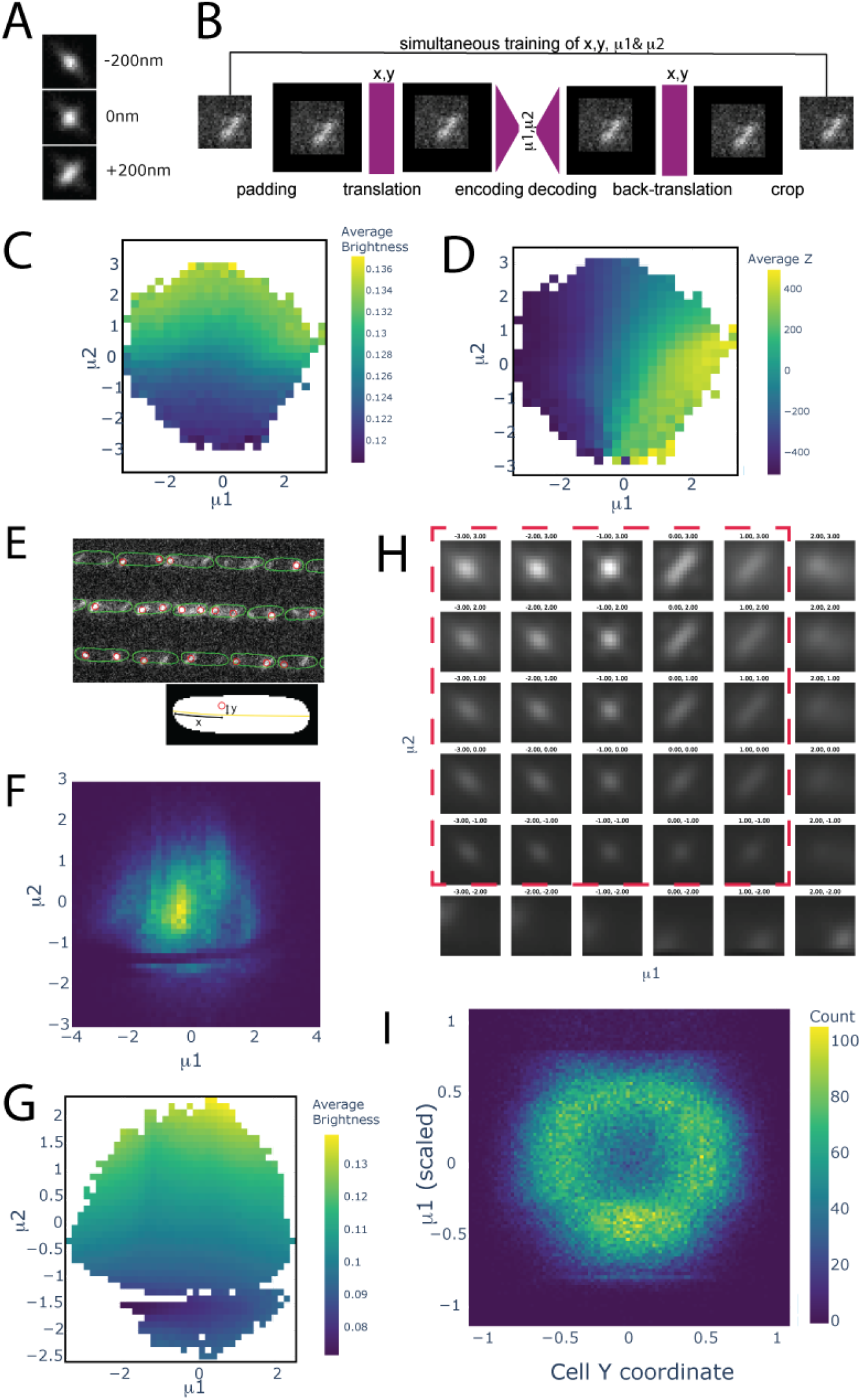
A. Three images of a bead at different z positions acquired through an astigmatic lens. B. In the autoencoder network the input image is encoded into two latent variables, such that the original image can be decoded from these two values alone. The x, y translation to center the emitter is handled by a spatial transformer network outside of the autoencoder. The weights in all networks (*purple*) are *trained* simultaneously to minimize the difference between the input and the output image. C.D Sample images from a spline-PSF model taken at different (known) z-values as represented in 2D latent space color-coded by the emitter brightness in C and the z position in D. E *top. E. coli* cells with fluorescent loci labels outlined with a segmentation mask from phase contrast. *bottom*. Localization of loci in the cell’s internal x and y coordinates. F. Distribution of chromosome locus images represented in 2D latent space color-coded by abundance. G. Function of chromosome locus images represented in 2D latent space color-coded by brightness. H. Decoded PSFs corresponding to different coordinates in latent space. I. Reconstruction of the yz distribution of chromosome loci.

In an autoencoder network, the input image is encoded into a low-dimensional “latent” space, such that the original image can be decoded as accurately as possible from the latent space information alone. We hypothesized that if we train an autoencoder network ^5^ with a narrow bottleneck to encode and decode astigmatic images of emitters at different, but unknown, z-positions, the latent space must include the z-information to be able to decode the image. Furthermore, if we center the emitter in the xy-plane (Fig 1B), we hypothesized that the most important information to encode a 3D-PSF in addition to the z-position should be the intensity. To test this hypothesis, we trained an autocoder network with only two latent variables on simulated images of point emitters at random z positions. To create a smooth mapping of properties from image space to latent space, we used a variational autoencoder which ensures that a small change in the input image only makes a small change in latent space. In practice, this is achieved by encoding images as a normal distribution in latent space and then sampling from this distribution. This pushes the network to learn representations where a small change in image space does not cause a large change in image space. To evaluate the trained network, we plotted the brightness and z positions of the emitters as a function of their latent space coordinates in Figs 1C and 1D, respectively. The brightness increases monotonically and smoothly with the second latent variable (µ2) and the z position increases similarly but with the first latent variable (µ1). The network has thus encoded brightness and z position as we hypothesized.

To evaluate if a similar encoding of the z position can be obtained on emitters in living cells, we imaged an *Escherichia coli* strain where a chromosomal locus was labeled with an array of 12 operator sequences bound by fluorescent transcription factors ^6^ (Fig 1E). This locus has previously been shown to be located mainly in the outer part of the nucleoid ^7^, giving rise to a characteristic crater-like 2D location localization distribution in a y-z cross-section of the rotationally symmetric cells. We now ask, can we use an autoencoder to obtain a similar distribution without explicit z-reference information?

To train the autoencoder we used small images cropped out around the chromosomal emitters that were first identified with a wavelet transform and then approximately located in x-y by Gaussian fitting. One million images were used for training. In Fig 1, we show the distribution of emitters in latent space. In this case, we did not have access to the ground truth z-location from the simulated data. However, the emitter brightness informs us which latent variable encodes the intensity information (Fig 1G) and the other variable should, by inference from the simulated data, encode the z-information. By plotting the decoded PSF-images corresponding to different coordinates in latent space (Fig 1H), we confirm that depth information is encoded in the first latent variable as the eccentricity of the PSF changes similar to the bead-stack (Fig 1A) along this dimension. In Fig H we also see that only the regions outlined by a red dashed square encode meaningful PSFs, since lower values in the second latent dimension imply very low brightness and higher values in the first latent dimension encode patterns corresponding to, for example, double emitters. For this reason, we only use data within the dashed region for downstream analysis. This also corresponds to 80% of emitters (Fig 1F).

The cells were segmented in x-y based on phase contrast imaging data ^8^, and the outlines were used to map the located emitters in the individual cells’ internal xy-coordinate system. When we make a scatter plot of the yz-coordinate system where z is the first latent variable, a crater-like distribution appears (Fig 1I). To obtain the absolute magnitude of the latent space variable, it was scaled such that the marginal distribution in z was the same as in y (Methods). Since the distribution of emitters excludes the center of the cell both in y and z, we conclude that the z-coordinate is accurately encoded in the first latent variable.

This exercise proves that the z information of individual emitters can be obtained directly from sample data without a reference model of the PFS. To obtain the correct scale when converting the latent variable to a z coordinate in image space, we used the additional information that the marginal distribution in y and z should be the same. Similarly, if it is previously known that the data is uniformly distributed in z, this information can be used to find the monotonous function of the latent variable that makes the marginal distribution of z uniform. When the marginal distribution in z is unknown, we suggest that training data be complemented by pairwise acquisition of the same emitter with a known z-shift of a few hundred nm. When the displaced emitter pairs are analyzed in latent space this results in a local scale bar for how a known change in z causes a change in latent space.

In summary, the proposed method for latent space encoding of z can be trained directly on the experimental data, without independent information about the PFS.

## Methods

### Network architecture

The network architecture combines a Beta-Variational Autoencoder, β**-**VAE ^9^ with a Spatial Transformer Network, STN ^10^. The β**-**VAE consists of mirrored encoder and decoder networks that consist of blocks of residual convolutional neural networks. The β refers to the β parameter that is used to weigh the importance of the Kullback-Leibler divergence (KLD) loss during training. The STN is constrained to translating the image in x and y. The original image is padded from 16×16 to 32×32 before being input into the STN. The STN outputs the image, translated through subpixel interpolation, and the coordinates describing the translation. The padded and translated image is input into the encoder of the VAE, which encodes the image into a 2D Gaussian in the 2D latent space. A sample from the encoded Gaussian is input into the decoder which outputs a 32×32 image. The image is then translated back in the opposite direction of the original translation by the STN. The translated image is cropped to 16×16 and output.

### Training

The entire network consisting of the STN and VAE is trained simultaneously via stochastic gradient descent using Adam ^11^. The loss function combines a reconstruction loss, binary cross-entropy, with a regularizing KLD loss. This loss regularizes the latent distribution by penalizing deviations from a gaussian prior. Without the KLD loss the model would only learn the parameters that are optimal for reconstructing the images. But we want the model to learn a smooth distribution that models the point spread function as a function of the Z-position. In practice these two losses have to be balanced in order to achieve this, the KLD loss is therefore scaled by a factor β, in this case 0.043. The images are encoded as 2D normal distributions with a predicted mean and variance in each dimension. The mean of the latent variable encoding the Z-position can be interpreted as a prediction of the emitter’s Z-position. However these predictions exhibit some undesirable behavior. For darker images the encoded distributions are centered close to the middle of the latent space with large variance. This makes it difficult to interpret the encodings as predictions of the emitter’s Z-position. To overcome this, the model training was divided into two phases. During the first phase, the entire network was trained as described above. This training ensures that the latent space is regularized. The expressive capabilities of the network are defined by the decoder, therefore we reasoned that a regularized decoder will effectively regularize the entire model, regardless of the behavior of the encoder. As a second training step, the weights of the pretrained decoder were frozen and the encoder and STN were reinitialized. Then we trained the encoder and STN to reconstruct the images using the frozen decoder with the regularizing loss term removed.

### Fluorescence microscopy imaging

All imaging was performed with a Ti-E (Nikon) microscope with a 100X immersion oil objective (Nikon, NA 1.45, CFI Plan Apochromat Lambda D MRD71970) for phase contrast and wide field epi-fluorescence microscopy. A Kinetix sCMOS camera (Teledyne Photometrics) was used for the fluorescence image acquisition, with the camera acquisition being triggered by a 515 nm laser (Fandango 150, Cobolt) for fluorophore excitation with a function generator (Tektronix). The same laser was used for fluorescent bead samples and live cell samples. The fluorescence images were acquired at 150 ms exposure time and 5 W/cm^2^ power density. The emitted fluorescence was passed through a FF444/521/608-Di01 (Semrock) triple-band dichroic mirror, a cylindrical lens, a BrightLine FF580-FDi02-T3 (Semrock) dichroic beamsplitter, and a BrightLine FF01-505/119-25 (Semrock) filter onto the sCMOS camera. The microscope was run using in-house plugins to MicroManager.

A DMK 38UX304 camera (The Imaging Source) was used to acquire phase contrast images. The exposure time was 50 ms and the light source used for phase contrast was a 480 nm LED and a TLED+ (Sutter Instruments). The light transmitted through the sample was passed through the same FF444/521/608-Di01 (Semrock) triple-band dichroic mirror as the fluorescence and reflected onto a different light path than the fluorescence using a Di02-R514 (Semrock) dichroic mirror. The light was directed onto the camera with a lens relay system, which included a phase ring at the back focal plane.

### Bead data acquisition

#### Acquisition of z-stacks for the PSF model

Fluorescent beads with a 100 nm diameter (TetraSpeck, Thermo Fisher) were diluted 1:50 in Milli-Q water and sonicated for 20-30 minutes, followed by application on an agarose pad sample. Z-stacks of fluorescent bead images were acquired with a custom BeanShell script in MicroManager.

#### Simulated training data

Z-values and photon counts were sampled from a uniform distribution and images of single emitters centered in the middle of the image were sampled from a spline-model^12^ that was fit to bead stacks. These images were corrupted with gaussian noise. A standard VAE with β=1 was trained on these samples.

### Bacterial data acquisition

#### Imaging

The *E. coli* strain imaged had the genotype MG1655 *rph+* Δ*malI::frt intC::P59-malI-SYFP2-frt gtrA::P58-mCherry2-parB-SpR 478731::KanR::MalOx12*, where 478731 indicates the genomic position of the locus label. Inoculation of the strain from a cryo-stock was done one day before the experiment into M9 minimal medium with 51 µg/ml Pluronic F-108 (Sigma-Aldrich 542342), 0.4 % succinate, and 1× RPMI 1640 amino acid solution (Sigma) at 30 °C. After overnight growth at 30 °C in a shaking incubator (200 rpm), the culture was diluted 1:100 into the same medium and loaded in a microfluidic chip ^13^. Phase contrast images were acquired every minute and fluorescence images were acquired every 2 min at 16 positions on the microfluidic chip sample over 10 hours of imaging, with each position including approximately 200 cells in the field-of-view.

#### Transformation between phase contrast and fluorescence cameras

As phase contrast and fluorescence images were acquired with two different cameras, a transformation matrix was fit using landmark-based registration of fluorescent bead images on both cameras. The sample of fluorescent beads with 500 nm diameter (TetraSpeck, Thermo Fisher) used for the registration was prepared in the same way as the 100 nm diameter beads, but with a 1:100 dilution before application on the agarose pad.

#### Internal cell coordinates

Segmentation of phase contrast images was performed with an Omnipose network ^14^ used to determine outlines of bacterial cells in the images. The fluorescent emitters were detected using the wavelet method ^15^ and the emitter coordinates were localized by 2D Normal distribution fitting of the emitters in the fluorescence image^16^. The emitter’s internal coordinates were estimated by fitting a cell backbone as a second-degree polynomial between the cell poles and estimating the distance to the cell poles for the long axis coordinate, and the signed distance to the cell backbone for the short axis coordinate.

Before estimating the emitter’s internal coordinates, the localized coordinates in the fluorescence images were transformed using the transformation matrix between fluorescence and phase contrast cameras.

## Acknowledgments

This work was made possible by grants from the ERC (advanced grant no. 885360), the Knut and Alice Wallenberg Foundation (2019.0439), and the eSSENCE e-science initiative. Author contributions: J.E. conceived autoencoding of z, A.T. designed networks and the training schemes, wrote the code and analyzed the data, K.G. performed the experiments and analyzed the data. All authors wrote the paper with input from the other authors. We are grateful for helpful discussion with Yoav Shechtman and comments on the manuscript from David Fange and Irmeli Barkefors.

